# Recallable but not Recognizable: The Influence of Semantic Priming in Recall Paradigms

**DOI:** 10.1101/2020.01.06.896795

**Authors:** Jason D. Ozubko, Lindsey Ann Sirianni, Fahad N. Ahmad, Colin M. MacLeod, Richard James Addante

## Abstract

When people can successfully recall a studied word, they should be able to recognize it as having been studied. In cued recall paradigms, however, participants sometimes correctly recall words in the presence of strong semantic cues but then fail to recognize those words as actually having been studied. Although the conditions necessary to produce this unusual effect are known, the underlying neural correlates have not been investigated. Across two experiments, involving both behavioral and electrophysiological methods (EEG), we investigated the cognitive and neural processes that underlie recognition failures. Experiments 1A and 1B showed that, in cued recall paradigms, presuming that recalled items can be recognized is a flawed assumption: Recognition failures occur in the presence of cues, regardless of whether those failures are measured. Experiment 2 showed that successfully recalled words that are recognized are driven by recollection at recall and by a combination of recollection and familiarity at recognition; in contrast, recognition failures are driven by semantic priming at recall and followed by negative-going ERP effects consistent with implicit processes such as repetition fluency and context familiarity at recognition. These results demonstrate that recall—long-characterized as predominantly reflecting recollection-based processing in episodic memory—can at times also be served by a confluence of implicit cognitive processes.

---

Dual-process models of episodic memory propose that memory consists of two distinct processes: familiarity and recollection (Gardiner, 1988; Jacoby, 1991; Mandler, 1980; Tulving, 1985; see Yonelinas, 2002, 2010 for reviews). Whereas familiarity reflects an intuitive feeling that a stimulus was recently experienced, recollection is considered to be a reconstructive process that retrieves the details and contextual information about an item’s prior occurrence. In recognition tests, because test probes are provided to participants, dual process models hold that either an intuitive sense of familiarity or an explicit conscious recollection can be used to judge whether the probe was recently experienced. For recall tests, however, because participants must actively retrieve an item from memory, conscious recollective processes are often deemed necessary because feelings of familiarity are seen as being unable to retrieve or produce items. In terms of memory paradigms, then, few tasks or behaviors are thought to be more explicit or demanding of recollection than recall.

By this standard account of recall, words that can be recalled should be easily recognized, as recalling a word entails recollective success, and recollection can be used to recognize words as previously studied. Indeed, it seems an intuitive contradiction for a participant to recall a word that they could not recognize, because of the assumption that if a memory representation is sufficiently strong to be recalled, surely it is sufficient to be recognized as well. Yet, in forced-recall-recognition procedures, where participants must produce words in response to cues and then recognize those items as either “old” (studied) or “new” (i.e., a guess) participants reliably recall studied words which they then cannot recognize as “old” (Allan & Rugg, 1998; Angel, Fay, Bouazzaoui, Granjon, & Isingrini, 2009; Angel, Fay, Bouazzaoui, & Isingrini, 2010; Angel, Isingrini, et al., 2010; Thomson & Tulving, 1970; Tulving & Osler, 1968; Tulving & Thomson, 1973). This unusual phenomenon is termed *recognition failure of recallable words*.^1^

For recognition failures to occur, acts of production and of recognition must occur. First, participants must produce a correct word and then they must fail to recognize it. Accounts of recognition failure, such as encoding specificity accounts, have generally focused on this second act—the processes that underlie the recognition failure step. These accounts focus on why words cannot be recognized, and how the semantic interpretation of words at study and test could cause recognition processes to fail (Thomson & Tulving, 1970; Tulving & Osler, 1968; Tulving & Thomson, 1973). For example, an encoding specificity account suggests that if the word GENERAL was encoded at study in the context of ‘*military*,’ a participant might imagine an army general commanding troops and may thus not recognize GENERAL at test if it is presented in the context of the cue ‘*specific.’* In essence, although the two words (GENERAL and GENERAL) are nominally identical, they have completely different meanings (“soldier” and “non-specific”). Thus, for all intents and purposes, GENERAL and GENERAL are *not* the same word and, from this semantic perspective, there is no reason to assume that a participant should recognize them as the same (see Martin, 1975, for more on this argument).

Although understanding the processes that lead to failure of recognition is important, such recognition failures cannot occur unless words that cannot be recognized are first produced. Yet surprisingly, despite decades of research, the mechanism that drives the production of recognition failures has been explored very litte. Accounts like the encoding specificity account suggest a mechanism through which recognition can fail for recalled words, but these accounts do not elucidate the processes that underlie the actual recall and generation of these items in the first place. Hence, no account has yet been put forward to explain why recognition failures are often *produced* at a level above what would be expected by free-association norms.

One implication of failing to consider the mechanisms that underlie the production of recognition failures is that recognition failures may regularly be contaminating measures of recall performance in more traditional recall paradigms where no explicit recognition decision is solicited from participants. Indeed, recognition failures can only be identified in forced-recall-recognition procedures when participants must recognize their own recalls. In paradigms that simply ask participants to recall words but that do not require a recognition decision, it is unknown how often recognition failures are occuring, as they would outwardly appear simply to be correct recalls. Hence, to date, it is also unclear how often recall results are biased by these recognition failures.

The present study sought to investigate recognition failures to address three important issues. Experiment 1 used behavioral methods to investigate the frequency of recognition failures in both free and cued recall: during both types of recall, we examined recall rates when a recognition decision was required and when it was not. It is possible that when recognition decisions are forced, participants adopt a more liberal threshold when recalling words, leading them to produce words that they cannot recognize but which they would never produce if they knew that a recognition decision was not required. Hence, recognition failures could be an artifact of forcing recognition, and may not be contaminating recall results in non-forced-recognition procedures. To preface, Experiment 1 ruled out this possibility and showed that forced-recognition procedures do not influence the production of recall responses.

Experiment 2 used neuroscientific methods of event-related potentials (ERP) to further investigate the cogntive processes that underlie the production of recognition failures— investigating recall itself in a way that has not been done before. Although it is common to view recall as a primarily recollective phenomenon, we will demonstrate that cued recall is a complex, multi-stage process of production and recognition that incorporates recollection, semantic priming, and familiarity.

## Experiments 1A & 1B

Recognition failures are an apparent paradox of recall—words that can be recalled but that cannot be recognized. However, in any recall paradigm that does not require a recognition response (i.e., most typical cued recall paradigms), it is unknown whether a given recalled word could indeed be recognized. Hence, the frequency of recognition failures occurring in standard free recall and cued recall procedures is unknown. While, as noted earlier, prior work has identified instances in which recognition failures exist in cued-recall-and-recognize paradigms, specifically with respect to free recall, no study to date has investigated the possibility that recognition failures exist.

Experiment 1 served two purposes. First, in Experiment 1A, we contrasted free recall and cued recall in forced-recognition procedures to investigate the frequency of recognition failures in each. Are recognition failures primarily a product of experimenter-provided explicit cues, or do they occur in free recall as well, where the only cues available are those implicitly generated by the participant? Second, we investigated whether the inclusion of the recognition decision itself might produce recognition failures. That is, to observe recognition failures, is it necessary to require participants to recall words and then to recognize their own recalls as either “old” or “new”?

Because this task might seem a bit odd to the participant, who may wonder why they have to recognize words that they are already recalling, this may induce participants to act in an artificial way: producing words more liberally during recall because they are fully aware that they can reject words at the recognition stage. To investigate this possibility, Experiment 1B replicated Experiment 1A with one key modification—that there was no post-recall recognition decision. In Experiment 1B’s design, participants were simply asked to recall words freely or in response to cues. If participants recalled fewer words in Experiment 1B compared to Experiment 1A, and especially if the number of words in Experiment 1B declined by the rate that recognition failures were seen to occur at in Experiment 1A, this would suggest that recognition failures may be an artefact of forced-recognition in recall paradigms. On the other hand, if the overall number of words recalled in Experiment 1B matched the number of words recalled in Experiment 1A, this would suggest that recognition failures are present in Experiment 1B but are going undetected and hence are being incorrectly categorized as true recalls.

## Method

### Participants

In total, 126 undergraduate students from the University of Waterloo participated in Experiments 1A and 1B for credit. In Experiment 1A, 30 students participated in the Free Recall condition and 31 participated in the Cued Recall condition. In Experiment 1B, 37 students participated in the Free Recall condition and 28 participated in the Cued Recall condition.

### Materials

A word pool of 200 cue-target pairs was created from the free association norms of Nelson, McEvoy, and Schreiber (2004). For present purposes, the backward association norms compiled by Nelson et al. were of principal interest. These norms are arranged by target words instead of cue words: For each target word, the norms provide a list of the cue words that gave rise to that target word together with the probability that each cue word gave rise to that particular target word. For example, if RIGHT was the target word of interest, Nelson et al. list *left*, *wrong*, *correct*, and *accurate* as cue words which gave rise to RIGHT during free association with probabilities of .93, .72, .23, and .16, respectively. Note that for our purposes, cases of repetition either between the target and cues or within the cues of different items were eliminated. For example, if PRINCESS was a target itself but *princess* was also a cue for the target KING, then one of these items was eliminated to avoid repetition. Additionally, if *universe* was a cue for the target word WORLD but also for the target word GALAXY, then one of these items was eliminated. We used the strongest associate of each target as its cue. On average, the normed probability that the strong associate cues would give rise to their respective target words was .58 (*SD* = .13). Stimuli were randomly selected for each participant.

### Procedure

In Experiment 1A, each participant studied 24 words randomly selected target words from the stimulus pool and was then tested either with 48 semantic associate test cues (in the Cued Recall condition) or with 48 un-cued trials (in the Free Recall condition). In the Cued Recall condition, for half of the test trials, strong semantic associate cues of the studied targets were used; for the other half of the test trials, strong semantic associate cues of unstudied words were randomly selected from the stimulus pool. These two types of trials were randomly inter-mixed at test and participants were explicitly informed that some cues would be more useful than others for retrieving studied words.

In the study phase, each of the 24 target words was presented individually in the center of the screen for 2 s with a 0.5 s inter-stimulus interval. At test, participants produced words by typing them at the keyboard, following each by the ENTER key. Participants were informed at test that, after producing a word, that word would re-appear in the center of the screen along with the labels “old” and “new” below it. Participants were told to press the M key to indicate that a word was old (i.e., studied) or the C key to indicate that a word was new (i.e., unstudied).

Experiment 1B was conducted similarly to Experiment 1A except that participants were told only that they were to recall words that they had studied; they were not required to recognize their responses. On each test trial, then, participants were asked to recall a word that they had studied by typing a response into the keyboard and pressing ENTER. Participants could skip trials (in the Cued Recall test) or end the recall session when they were finished (in the Free Recall test). Thus, there was no forced-response component to Experiment 1B.

### Results

The mean numbers of words produced in Experiments 1A and 1B can be seen in Figure 1. Regarding overall recognition accuracy, in Experiment 1A, mean hit rates (probability of identifying an old word as “old”) and miss rates (probability of identifying an old word as “new”) of recalled words were .78 and .22 respectively (<u>SE</u> = .02) in the Cued Recall condition and .93 and .07 respectively (<u>SE</u> = .03) in the Free Recall condition. Mean correct rejection rates (probability of identifying a new word as “new”) and false alarm rates (probability of identifying a new word as “old”) were .87 and .13 respectively (<u>SE</u> = .02) in the Cued Recall condition and .94 and .06 respectively (<u>SE</u> = .01) in the Free Recall condition.

**Figure 1.**
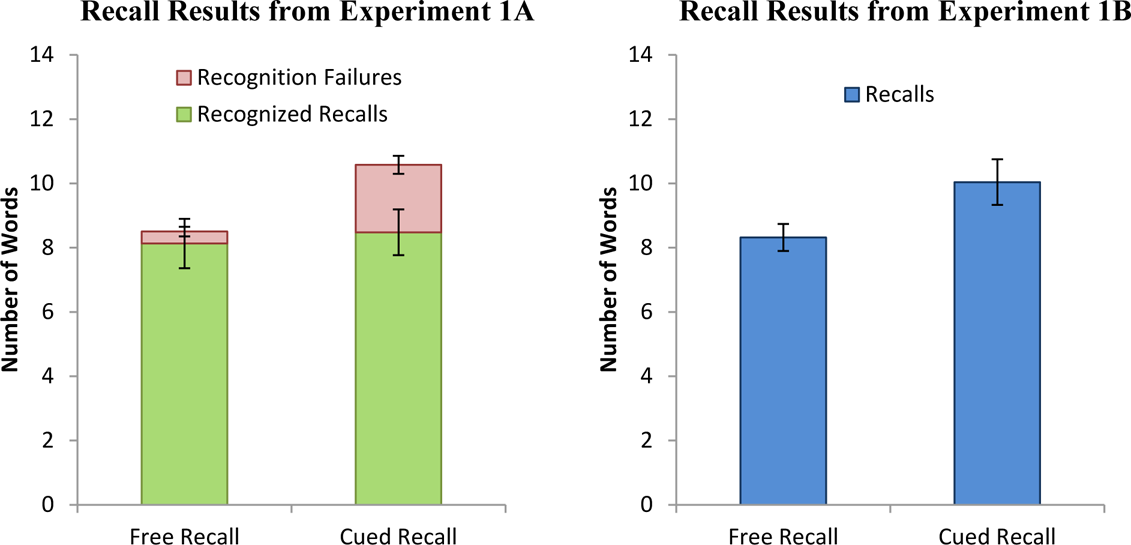
Mean number of words recalled in Experiments 1A and 1B. In Experiment 1A, the mean number of words recalled is separated based on whether the words were recognized as old or not recognized (i.e., recognition failures). Error bars are plotted separately for mean number of recognized recalls and mean number of recognition failures respectively at the top of each bar. In Experiment 1B, because there was no recognition phase, the mean number of words recalled is all that can be reported. Error bars represent the standard error of the mean in all cases.

In Experiment 1A, the number of recognized recalls in the Cued Recall and the Free Recall conditions did not differ significantly, t(59) = 0.74, <u>p</u> = .34, <u>d</u> = 0.09. There were, however, significantly more recognition failures produced in Cued Recall than in Free Recall, t(59) = 5.41, <u>p</u> < .01, <u>d</u> = 1.40, and thus there were significantly more old items produced in general in Cued Recall than in Free Recall, t(59) = 2.14, <u>p</u> < .05, <u>d</u> = 0.54. Significant recognition failures were observed only in the Cued Recall condition.

In Experiment 1B, more old words were correctly recalled in Cued Recall than in Free Recall, t(63) = 2.18, <u>p</u> < .05, <u>d</u> = 0.55. Importantly, there was no difference in the number of old words recalled in the Cued Recall conditions of Experiments 1A and 1B, t(57) = 0.56, <u>p</u> = .58, <u>d</u> = 0.14. As well, there was no difference in the number of old words recalled in the Free Recall conditions of Experiments 1A and 1B, t(65) = 0.22, <u>p</u> = .82, <u>d</u> = 0.06. Hence, the absence of the recognition decision did not affect the overall recall rates in Experiment 1B compared to Experiment 1A. Consequently, the recognition failures observed in Experiment 1A must also have been present in Experiment 1B but were undetected.

## Discussion

Experiment 1 sought to address two issues. First was the question of whether recognition failures are present both in cued recall and in free recall, or whether they are present only in cued recall. Experiment 1A showed clearly that recognition failures are virtually absent from free recall yet readily occur in cued recall. Hence, the presence of cues appears to be critical for the production of recognition failures. Second was the question of whether forcing participants to recognize their recalls results in recognition failures. Experiment 1B found the same overall rate of cued recall as Experiment 1A, suggesting that the recognition failures that were identified in Experiment 1A were also present in Experiment 1B, where no recognition decision was present. The clear implication is that some portion of recalled words in Experiment 1B were actually recognition failures but these could not be detected given that the canonical recall procedure lacked a recognition decision.

Having now demonstrated that recognition failures can easily be induced in cued recall, the obvious follow-up question is why: What factors lead to the production of recognition failures in the first place? To investigate this issue, we turn to an electrophysiological approach to examine the ERP signatures of memory.

### Experiment 2

Although traditional accounts provide some explanation for why recognition failures are not identified (Thomson & Tulving, 1970; Tulving & Osler, 1968; Tulving & Thomson, 1973; Tulving & Watkins, 1977; for a review, see Gardiner & Nilsson, 1993), these accounts do not elucidate the processes that underlie the *production* of recognition failures. Indeed, traditional accounts suggest that recognition failures are simply “happy accidents.” The idea is that, on trials in which the participant is unable to recall the correct studied word, they simply guess. When guessing, they should be no more likely to produce the correct studied word to a related cue than they should be to produce the “correct” target to a cue related to an unstudied item. Yet recognition failures are often produced to cues more frequently than would be expected by chance, suggesting that some form of fluency (Leynes & Zish, 2012; Leynes & Addante, 2016), explicit familiarity, or perhaps implicit priming must be driving these responses. Unfortunately, neither traditional accounts nor existing behavioral data provide much insight into these underlying processes. We therefore examined ERP correlates of these processes to provide a more thorough investigation of the processes that drive the production and subsequent recognition of recognition failures.

Our main motivation for using ERPs was the ability of ERP techniques to characterize distinct mnemonic processes using well-established electrophysiological markers. By using ERPs, our goal is to identify the mnemonic processes that underlie both the production and the recognition of words in cued recall, and to provide a more sophisticated analysis of the processes that give rise to recognition failures than has been reported in the past. Indeed, although ERPs have commonly been applied to recognition, they have rarely been applied to recall since it has traditionally been difficult to link precise time stamps of events to the physiology recorded; thus, another novel aspect of this work is that it will be one of the few, thorough, ERP examinations of mnemonic processing during recall generally. Before proceeding to our experiment, we describe the mnemonic processes that ERPs will allow us to characterize in more detail via the ERP correlates of recognition and recall, respectively.

### ERP Correlates of Recognition

ERPs have been widely studied in recognition memory and researchers have been able to characterize distinct spatio-temporal waveforms that can be used as reliable markers of retrieval processes (Addante, Ranganath, & Yonelinas, 2012; Duzel et al., 1997; Friedman & Johnson, 2000; Rugg & Curran, 2007; Rugg, Mark, et al., 1998). Generally, ERPs elicited during recognition tests have been shown to differentiate whether an item has been previously studied (Addante, Ranganath, & Yonelinas, 2012). At a more process level, ERPs also have been used to differentiate specific retrieval processes during memory tests.

Correctly recognized studied items show an increased positivity compared to correctly rejected new items, a finding dubbed the “old/new effect” (Allan, Doyle, & Rugg, 1996; Allan, Wilding, & Rugg, 1998). This general old/new effect is comprised of at least two temporally, topographically, and functionally distinct components that have been shown to be correlates of explicit memory (i.e., recollection and familiarity) and of implicit memory (i.e., priming).

Whereas familiarity is often associated with an old/new difference that onsets relatively early after stimulus onset (∼300-500 ms) with a mid-frontal scalp distribution (referred to as a “mid-frontal old-new effect” or “FN400”), recollection is often associated with an old/new difference that onsets later (∼500-800 ms) and usually with a left parietal distribution, referred to as a parietal old/new effect or “LPC” (Left Parietal Component; for reviews, see Friedman & Johnson, 2000; Rugg & Curran, 2007).

In addition to characterizing explicit memory processes, ERPs are also useful in measuring implicit memory processes. Old/new ERPs associated with priming have been observed to onset early (∼300-500 ms), like familiarity, but in contrast are maximally distributed in more posterior scalp regions (Addante, 2015, Addante, Ranganath, Olichney, & Yonelinas, 2012; Addante, Ranganath, & Yonelinas, 2012; Bridger et al., 2012; Li, Mao, Wang, & Guo, 2017; Rugg, Mark, et al., 1998; Yu & Rugg, 2010; Bader & Mecklinger, 2017; although see Voss et al., 2012, and Mecklinger, Frings, & Rosburg, 2012, for differing discussion of these effects^2^). This final point is critical, because behavioral techniques for separating recollection and familiarity do not typically account for implicit memory processes. Indeed, several researchers have recently argued that implicit memory processes may often be mislabeled as familiarity effects, and thereby may distort interpretations of the data (Voss et al., 2012). ERPs offer an additional important benefit in that they potentially allow us to separate explicit from implicit memory processes (Addante, 2015; Mecklinger et al., 2012; Bader et al., 2012; Yu & Rugg, 2010; Rugg et al., 1998).

### ERP Correlates of Recall

Although ERPs have been examined extensively in recognition, they have been under-studied in recall. To our knowledge, no ERP studies have investigated cued recall using semantic associates as cues. However, using word-stem cues in forced-recall-recognition paradigms, several studies have documented reliable old/new ERP differences between hits and correct rejections (Allan et al., 1998; Angel et al., 2009; Angel, Isingrini, et al., 2010; Rugg, Mark, et al., 1998), allowing us to draw some inferences and predictions about recall. Nevertheless, because ERP studies of recall remain limited, our understanding of recall certainly would benefit from further in-depth investigation of the neural processes underlying recall.

Studies that have examined ERPs in recall and that have used designs appropriate for comparison with recognition have reported results consistent with the recognition studies of recollection-related and familiarity-related ERP effects. Although, the time windows associated with familiarity and recollection tend to occur approximately 200 ms later in recall than in recognition^3^. For example, in their examination of cued recall and source memory, Allan and Rugg (1998) demonstrated that the cued recall old/new effect is composed of a mid-frontal component which onsets 400-700 ms post-stimulus, and a left parietal effect which onsets 800-1200 ms post-stimulus. The posterior effect was associated with the amount of contextual detail retrieved for a given item, which is consistent with the neural correlates of recollection observed for recognition tasks. The earlier anterior effect was also associated with successful retrieval but not with the amount of contextual detail, consistent with the interpretation that it reflects familiarity-based processes.

Similarly, consistent with the recognition correlates of recollection and familiarity, Fay, Isingrini, Ragot, and Pouthas (2005) demonstrated that an early frontal effect, which onset 400-800 ms post-stimulus, was observed for both shallowly encoded and deeply encoded words in cued recall, but only the deeply encoded words demonstrated a late parietal effect, which onset 800-1100 ms post-stimulus. These convergent recall results suggest that despite both types of items demonstrating a familiarity component, only those that were deeply encoded showed a recollection component, corresponding with the ERP effects in recognition reported for shallow and deep encoding by Rugg and colleagues (Rugg, Walla, et al., 1998). Thus, ERP results observed in recall appear to closely parallel those observed in recognition, albeit with familiarity and recollection effects onsetting slightly later (by ∼200 ms) in recall than in recognition.

With regard to ERP patterns during recognition failures (misses) in recall, there is only one published result that provides any relevant analysis. Allan and colleagues (1996) reported that in a word-stem-cued forced-recall-recognition design, although recognized recalls (hits) showed more anterior positivity than misses or correct rejections (consistent with familiarity occurring for hits but not for misses), misses and correct rejections did not differ. This result could be taken to suggest that no process differentiates misses from correct rejections (a contrast typically used to reveal implicit memory differences). It should be noted, however, that Allan et al. did not examine early ERP effects (i.e., < 500 ms after stimulus onset) and therefore their results cannot speak to the possibility that recognition failures may be driven primarily by implicit priming processes that occur early. We thus turn to our own EEG investigation to examine the neural correlates that lead to both the production and identification of recognition failures in cued recall paradigms.

## Method

### Participants

Forty right-handed undergraduate students from California State University, San Bernardino were recruited to participate in exchange for monetary compensation of $10/hr. Participants were identified through screening processes as normatively healthy, free from any neurological disorders, fluent English speaking, and right-handed. Handedness was assessed via the Edinburgh handedness inventory (Oldfield, 1971), and other demographic criteria were established via self-reporting questionnaires.

### Materials

Experiment 2 used the same stimuli and word pool as Experiment 1; however, to obtain more observations of recalls and recognition failures per participant, more study and test trials were required per participant (described in Procedure) than were used in Experiment 1. To estimate the mnemonic procedures underlying both the production and recognition of recognition failures, ERP techniques require having sufficient trials for each participant in each of the respective conditions being compared so as to overcome inherently noisy signal-to-noise ratios for scalp-based electrophysiological analyses.

EEG was recorded during the test phase using a 32-channel EEG system (Brain Vision’s ActiCHamp design http://www.brainvision.com/actichamp.html) of Ag-CL electrodes un-referenced, at 500 Hz sample rate. This montage includes pre-amplifiers built into each electrode and electrically-shielded cabling. EEG sites were prepared and abraded with saline gel to facilitate optimal signal to noise connections with scalp sites in accord with the international 10-20 system (Klem, Luders, Jasper, & Elger, 1999). The electrode sites on the cap were filled with saline gel prior to insertion of the active electrodes. After insertion of active electrodes, impedance was reduced via gentle abrading of each site. Participants were instructed to minimize muscle tension, eye movements, and blinking during the test sessions. Electrooculogram (EOG) was monitored in the horizontal (lateral to each eye) and vertical (below and above the left eye) directions to eliminate trials contaminated by blink or eye-movement artifacts.

An SV-1 Voice Key (https://www.cedrus.com/sv1/) was used for logging precise voice responses during EEG recording of recall (see Procedure, Figures 2, 3). The SV-1 is a 100% digital device powered by an 18 MHz microprocessor, and is designed specifically for psychological experiments requiring a vocal response; it monitored the participant’s voice level at all times during the retrieval phase of the experiment. When the voice level rose above a user-specified threshold, the device transmitted that as a digital signal to the computer recording the EEG timestamps and behavioral data logs (see Figure 2).

**Figure 2.**
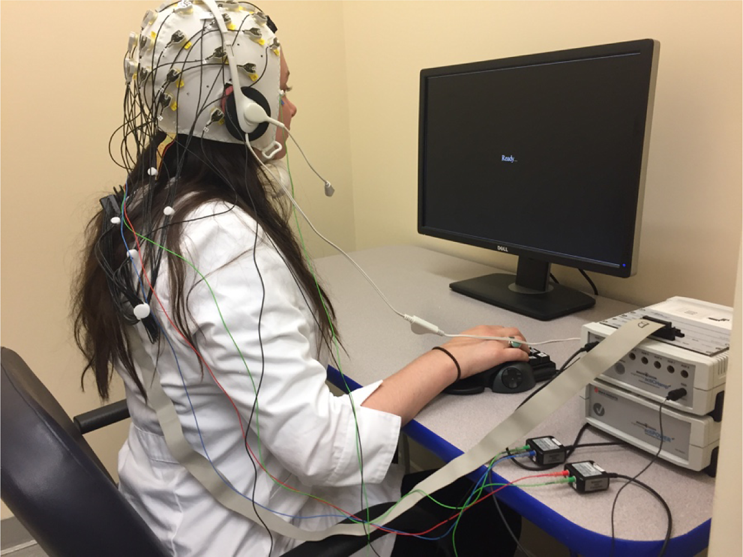
Example of the S-1 Voice Key device used to collect digitized time stamps of response times of cued recall for semantic associates in the current study, concurrent with EEG recordings.

### Procedure

The experimental procedure is outlined in Figure 3. The experiment consisted of 144 words during the study phase, broken down into 6 study blocks each containing 24 words. The test phase consisted of 288 words, divided into 6 test blocks each containing 48 words. Hence, half of the cues presented in each test session were semantic associate cues for the previously studied words and the other half were new cues for previously unstudied words. These two types of trials were randomly inter-mixed at test and participants were explicitly informed that some cues would be more useful than others for retrieving studied words. Stimuli for both the study and test phases were randomly selected for each participant. Instructions on task performance were read from a prepared script and reminders were given periodically. Short practice runs were used to ensure that instructions were understood and that participants were responding correctly during both study and test.

**Figure 3.**
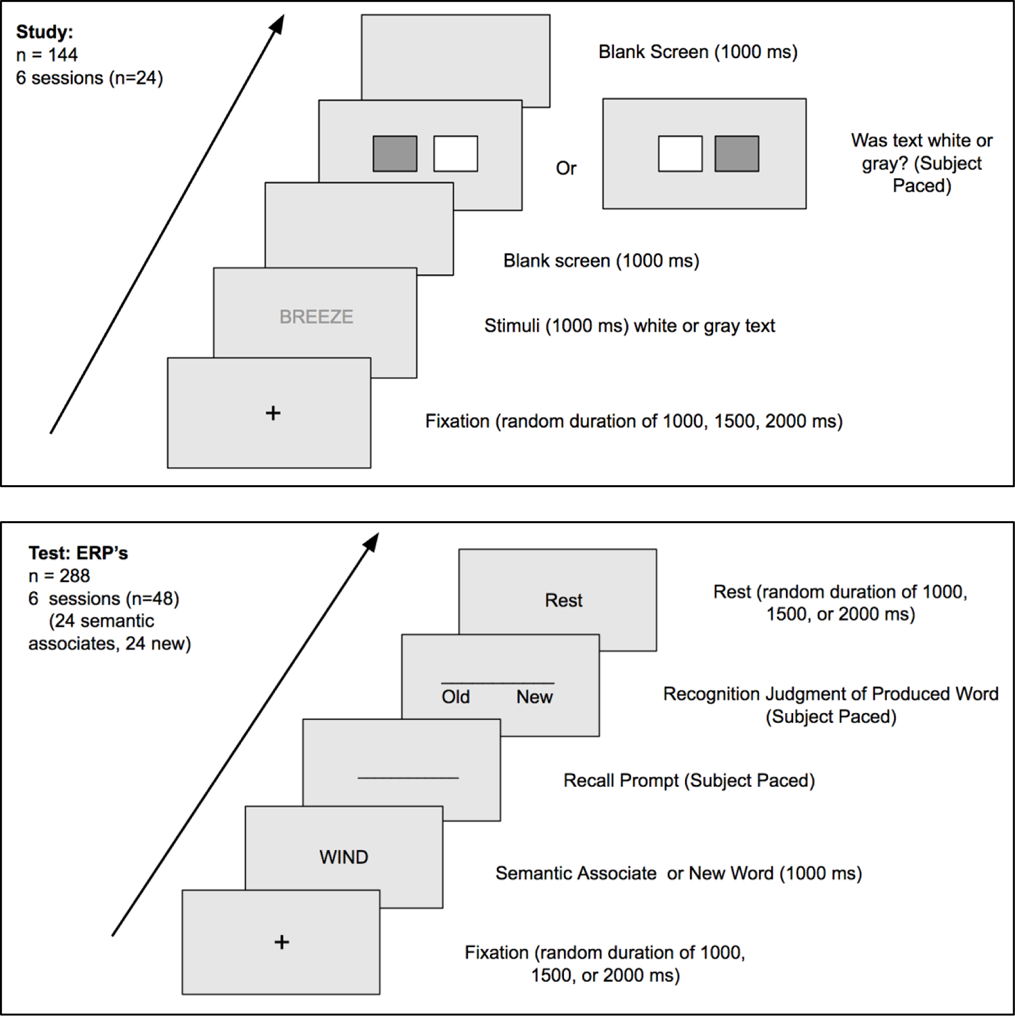
Experimental paradigm of Experiment 2. Top: The study phase (encoding). A total of 144 words, divided into 6 blocks of 24 words, were presented one at a time. Participants were instructed to select the color of the word, represented by gray and white boxes that alternated positions on the screen. Bottom: The test phase (retrieval). Two-hundred and eighty-eight new words, split into 6 blocks of 48 words, were presented one at a time, followed by a recall prompt. Half of the words were semantic associates of studied words and the other half were semantic associates of unstudied words, which we treated as ‘new’ words. Participants were prompted to recall the first word from the study session that came to mind, and then to recognize that word as “old” (from the study session) or as “new” (not from the study session).

In the study phase, participants first encoded a word presented on the screen for 1s and then were asked to indicate whether the font-color of the presented word was white or grey; they did so by pressing a response button corresponding to the location of grey and white boxes on the screen. The study was specifically designed to increase signal-to-noise ratio in ERP analyses for the otherwise relatively uncommon phenomenon of recognition misses of recalled words. Hence, the purpose of the perceptual encoding task was to engender a low-level of encoding that would increase the number of forgotten trials during the later recognition phase of the study. As a perceptual distractor specifically intended to not facilitate encoding, these boxes randomly alternated order, while the response keys “Grey” and “White” remained in the same location.

The retrieval test began with a fixation cross which appeared for a jittered duration of 1000, 1500, or 2000 ms. Next, a semantic associate either of a studied word or of a new word was presented on screen for 1 s. The recall prompt screen followed immediately. Participants were instructed to think of a studied word in response to the test cue or, if a studied word did not come to mind, to think of any new word. Participants were instructed to speak the recalled word aloud as soon as it came to mind. The voice key device digitally recorded the response time and integrated this event code into the EEG data. After their verbal response, participants were then prompted with an old-new recognition task and asked to identify the word that they had just produced as either “old” (from the study session) or “new” (not from the study session). To avoid introducing noise from eye-blinks into the neural data, participants were instructed not to blink when probes were on the screen; rather, they were to blink only during the ‘Rest’ screen (Addante et al., 2011).

*Electrophysiological Procedures and Analyses*. EEG data were analyzed using EEGLab (Delorme & Makeig, 2004) and ERP Lab analysis tool-boxes (Lopez-Calderon & Luck, 2014) for Matlab software. EEG data were re-referenced offline to the average of the left and right mastoid electrodes, then baseline corrected to the average activity 200 ms pre-stimulus by a polynomial detrending function of zero using a .1 Hz high pass filter, and down sampled to 256 Hz. The data was then epoched beginning 200 ms pre-stimulus presentation and continuing through 1800 ms post-stimulus presentation. This corresponded to the entire duration of each cue’s presentation to the participant and was categorized for analysis based on the subsequent responses given for recall and recognition. Independent components analysis (ICA) was performed using InfoMax techniques in EEGLab (Bell & Sejnowski, 1995) to accomplish artifact correction and the resulting data were individually inspected for artifacts, rejecting trials for eye blinks and other aberrant electrode activity. During ERP averaging, trials exceeding ERP amplitudes of +/-250 mV were excluded. Additional filtering, such as a 30 hz low pass filter, was applied to group ERPs to make figures correspond to the similar ‘smoothing’ function that the standard process of taking the mean voltage between a given two latencies accomplishes during statistical analyses of results (e.g., Addante, 2015).

Using the ERPLAB toolbox (Lopez-Calderon & Luck, 2014), automatic artifact detection for epoched data was also used to identify trials exceeding specified voltages, in a series of sequential steps as noted below. Simple Voltage Threshold identified and removed any voltage below −100ms. The Step-Like Artifact function identified and removed changes of voltage exceeding a specified voltage (100 uV in this case) within a specified window (200 ms), which are characteristic of blinks and saccades. The Moving Window Peak-to-Peak function is commonly used to identify blinks by finding the difference in amplitude between the most negative and most positive points in the defined window (200 ms) and comparing that difference to a specified criterion (100 uV). The Blocking and Flatline function identified periods in which the voltage did not change amplitude within a specified window adjusted for each subject’s trials (for reference see https://github.com/lucklab/erplab/wiki/Artifact-Detection-in-Epoched-Data, Lopez-Calderon & Luck, 2014). An automatic blink analysis, Blink Rejection (alpha version), used a normalized cross-covariance threshold of 0.7 and a blink width of 400 ms to identify and remove blinks (Luck, 2014).

For statistical analysis, we computed the mean amplitude of the ERPs across designated time windows at each electrode site for each participant and condition, and then assessed for reliable differences between the average of each respective condition. As described in introducing Experiment 2, the time windows associated with familiarity and recollection tend to occur slightly later (approximately 200 ms) in recall than in recognition because in cued recall tests participants are provided with a cue at the start of each test trial and must take a moment to generate their own candidate for recognition (see Generate-Recognize models of recall, e.g., Haist, Shimamura, & Squire, 1992; Jacoby & Hollingshead, 1990; Nobel & Shiffrin, 2001; Slamecka, 1972). Due to this more demanding nature of recall, the time windows that we identified for familiarity and recollection are approximately 300 ms later than those identified in other studies using different retrieval tasks.

For the familiarity contrast, we focused on the 600-900 ms time period at mid-frontal electrode sites, whereas for the recollection contrast we focused on the 900-1100 ms time window at parietal electrode sites. These time windows and electrode sites were selected a priori based on other studies of familiarity and recollection that identify time windows of 300-500 ms and 600-800 ms, respectively for each (Addante, Ranganath, Olichney, et al., 2012; Addante, Ranganath, & Yonelinas, 2012; Leynes, Landau, Walker, & Addante, 2005; Rugg & Curran, 2007). Implicit memory effects were assessed by creating a posterior electrode cluster of parietal and occipital electrodes during the 300-500 ms time window, consistent with the characterization of implicit memory effects in prior studies (Addante, 2015; Bader & Mecklinger, 2017; Bridger et al., 2012; Li, Mao et al., 2017; Li, Taylor, et al., 2017; Mecklinger et al., 2012; Rugg, Fletcher, et al., 1998; Strozak, Abedzadeh, & Curran, 2016; Voss et al., 2012; Voss & Paller, 2007, 2017; Yu & Rugg, 2010). Direct contrasts were assessed using corrected t-tests to assess differences between memory conditions.

ERP results are presented for each electrode region in temporal sequence through the epochs identified from our hypotheses based upon the existing literature (see ERP Correlates of Recall and ERP Correlates of Recognition), starting with earliest latencies (100-300 ms) and progressing through each subsequent period (300-500 ms, 600-900 ms, 900-1100 ms). We examined correctly recognized recalls, recognition failures, and correct rejections. For each time period, ERP effects are presented in order of our conditions of interest: recognized recalls and recognition failures, with each contrasted against correct rejections. Paired 2-tailed t-tests were used to assess conditions for each electrode cluster of regions during the a-priori defined latencies.

Electrode clusters were created for each hemisphere and region, based upon the international 10-20 system (Klem et al., 1999). The left frontal cluster included sites F3, F7 and FC5; mid frontal included sites Fz, FC1 and FC2; and the right frontal cluster comprised sites F4, F8 and FC6. Accordingly, the left parietal cluster included sites CP5, P3 and P7; mid parietal included Pz, CP1 and CP2; and the right parietal cluster comprised CP6, P4 and P8.

To maintain sufficient signal-to-noise ratio (SNR), all comparisons relied upon including the data of only those participants who met a criterion of having a minimum number of 12 artifact-free ERP trials per condition being contrasted (Addante, Ranganath, & Yonelinas, 2012; Gruber & Otten, 2010; Kim et al., 2009; Otten et al., 2006; cf. Luck 2016). For our main analysis, this yielded a sample of 28 participants; for our subsequent ERP analyses, the samples that met this criterion are indicated in each of those sections, respectively.

*ERP Conditions Analyzed.* Traditional approaches in extant research on cued recall for semantic associates has collapsed words produced from semantic associate cues and non-associate cues together into the same conditions, counting items as successfully recalled regardless of which cue initiated their retrieval, a frequently used practice in the recall literature (e.g.; Blaxton, 1989; Humphreys & Galbraith, 1975; Thomson & Tulving, 1970; Tulving & Osler, 968). However, by not specifying conditions based on whether responses were generated from semantic associate cues or non-associate cues, one may conflate processes if in fact distinct processes are used to arrive at these recalled items. We reasoned that it could be possible to gain a more sensitive measure of our conditions of interest if we used an approach that instead separated the conditions based on whether a word was produced in response to a semantic associate cue or to a non-associate cue.

Extant research on cued recall of semantic associates has typically defined recognized recalls (“hits”) as any instance in which an old word was produced at recall and later recognized as old, regardless of whether a semantic associate cue was given to the participant to induce it. By this definition of recognized recalls, studies collapsed across semantic associate and non-semantic associate conditions, so there was no distinction as to whether the participant produced merely ‘any’ old word from the study phase or they produced specifically the target old word of the semantic pairing (i.e., the participant produced “Animal” instead of “Stripe” to the cue word “Zebra”; although “Animal” was a studied word, “Stripe” was the target word for the cue “Zebra”). Accordingly, researchers have often defined recognition failures (“misses”) for these kinds of recall + recognition paradigms as instances in which an old word was produced at recall and the participant incorrectly identified the word as “new” for the recognition judgment. These, too, were not separated by whether a semantic associate or a non-associate cue was provided.

Likewise, extant research has also defined correct rejections as instances in which new words were produced at recall in response to either a semantic associate or a non-associate cue and then the participant also correctly identified the word as being new at recognition.

We reasoned that this approach was a good start for preliminary analysis but could also potentially be obscuring certain neurophysiological effects in ERPs because of the condition’s inherent coarseness resulting from collapsing across disparate conditions. Thus, we sought to create a more specified and targeted analysis. Therefore, we focused on the more specified criteria of semantic associate trials comprising conditions of recognized recalls and recognition misses. For our analyses, only recognized recalls and recognition misses that resulted from semantic associate cues, and correct rejections from non-associate cues, were analyzed. “Recognized recalls” were defined as instances in which participants *produced the target old word in response to a semantic associate cue in recall*, and then also went on to successfully rate the word that they had just produced as being ‘old.’ On the other hand, “recognition failures” were defined as instances in which participants successfully *produced the target old word during recall in response to a semantic associate cue* and then misidentified the word that they had just produced as a ‘new’ word (recognition failures of recalled items). “Correct rejections” were accordingly defined as new words *produced in response to non-associate cues*, which were then correctly identified as new words^4^. Only recognized recalls and recognition failures that were produced from semantic associate cues were analyzed; those resulting from non-associate cues were excluded in analyses.

## Results

### Behavioral Results

Behavioral data were assessed for accuracy and response time of participants’ responses on the memory test. Outlier participants performing beyond three standard deviations of the mean N=5) and those who also performed beneath chance level performance on the memory test (negative scores on accuracy, N = 6) were excluded from analysis; this resulted in the exclusion of the data of 11 participants, leaving 29 included subject data sets. The mean number of words produced in Experiments 2 can be seen in Figure 4A. Regarding recall production rates, of the 288 test trials per participant, participants produced a valid response (a coherent, non-repeated word) on a mean of 236 trials (SE = 3.47). This included a mean of 68 (SE = 2.64) old items produced in response to their matching cues^5^ and 163 (SE = 3.15) new items produced. Of new items produced, there were an average of 80 (SE = 3.85) correct rejections.

**Figure 4.**
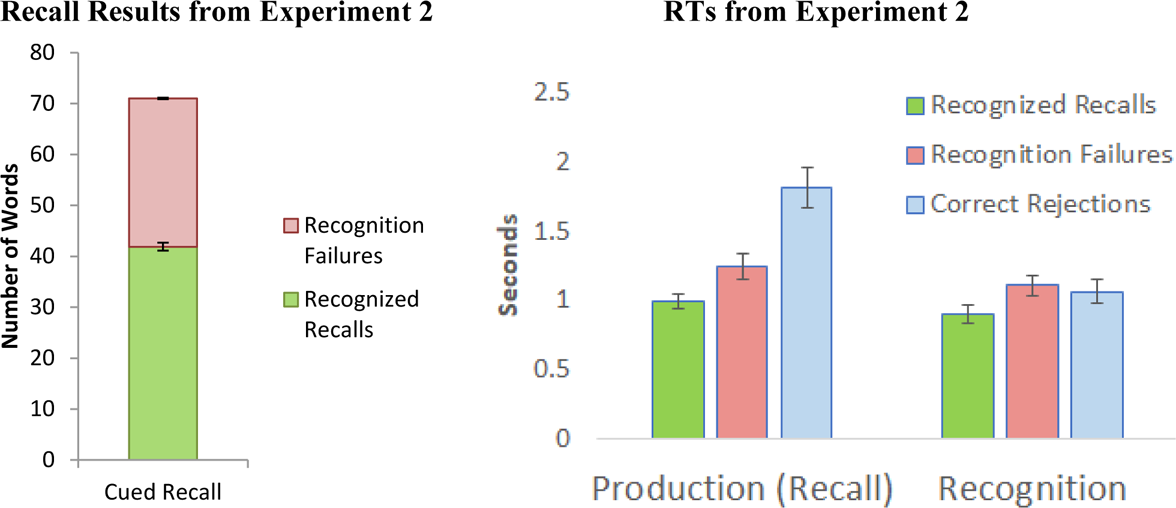
Mean number of words recalled in (Figure 4A) and mean RT for recall and recognition (Figure 4B) in Experiment 2. In Figure 4A, the mean number of words recalled is separated based on whether the words were recognized as old or not recognized (i.e., recognition failures). Error bars are plotted separately for mean number of recognized recalls and mean number of recognition failures respectively at the top of each bar. In Figure 4B, the mean RT to produce/recall a word and the mean RT to recognize a produced word is plotted separately for recognized recalls, recognition failures, and correct rejections. Error bars represent standard error of the mean in all cases.

Regarding overall recognition accuracy, mean hit rates and miss rates of produced words were .57 (SE = .03) and .43 respectively (SE = .03). Mean correct rejection and false alarm rates were .70 and .30, respectively (SE = .03 for both). Of the 68 words correctly recalled to their cues, participants produced a mean of 40 (SE = 2.73) recognized recalls and a mean of 28 (SE = 1.80) recognition failures of successfully recalled words. (Figure 4A).

In addition to overall recall and recognition rates, we also examined response time (RT) in Experiment 2. This was possible because Experiment 2 precisely measured recall via a digital voice response detector. Recognition RTs were measured as the time taken to produce a manual button press. Examining RTs for the recalled items could potentially shed light on the processes used for producing an old or new word in recall.

Mean RTs are shown in Figure 4B. The production of old items that would become recognized recalls was significantly faster than the production of both old items that would become recognition failures, t(28) = 4.24, <u>p</u> < .001, <u>d</u> = .60, and new words that would become correct rejections, t(28) = 7.75, <u>p</u> < .001, <u>d</u> = 1.39. The production of old items that would become recognition failures was also faster than that of new items that would become correct rejections, t(28) = 9.16, <u>p</u> < .001, <u>d</u> = .876. For recognition decisions, the mean RT of hits was significantly faster than those of misses, t(28) = 4.04, <u>p</u> < .001, <u>d</u> = .32, and correct rejections, t(28) = 4.79, <u>p</u> < .001, <u>d</u> = .412. Mean RT for misses did not differ significantly from that for correct rejections, t(28) = 1.70, <u>p</u> = .099, <u>d</u> = .104. Hence, in terms of both production and recognition times, recognition failures were slower than recognized recalls, but they were also distinct from correct rejections.

### Electrophysiological Results

#### ERP Results: Recall

We examined the ERP patterns for old words that were recalled and then went on to become either recognized recalls or recognition failures; we also assessed new words that went on to be correctly rejected. In the 100-300 ms latency, there were no reliable ERP effects between any conditions, as shown in Figure 5. In the 300-500 ms latency, recognized recalls (M = .49, SD = 2.38) were reliably less positive than correct rejections (M = .85, SD = 2.20) at right parietal sites, t(27 = 2.27, <u>p</u> < .05. Recognized recalls did not reliably differ from recognition failures nor did recognition failures differ from correct rejections during this latency for any electrode regions.

**Figure 5:**
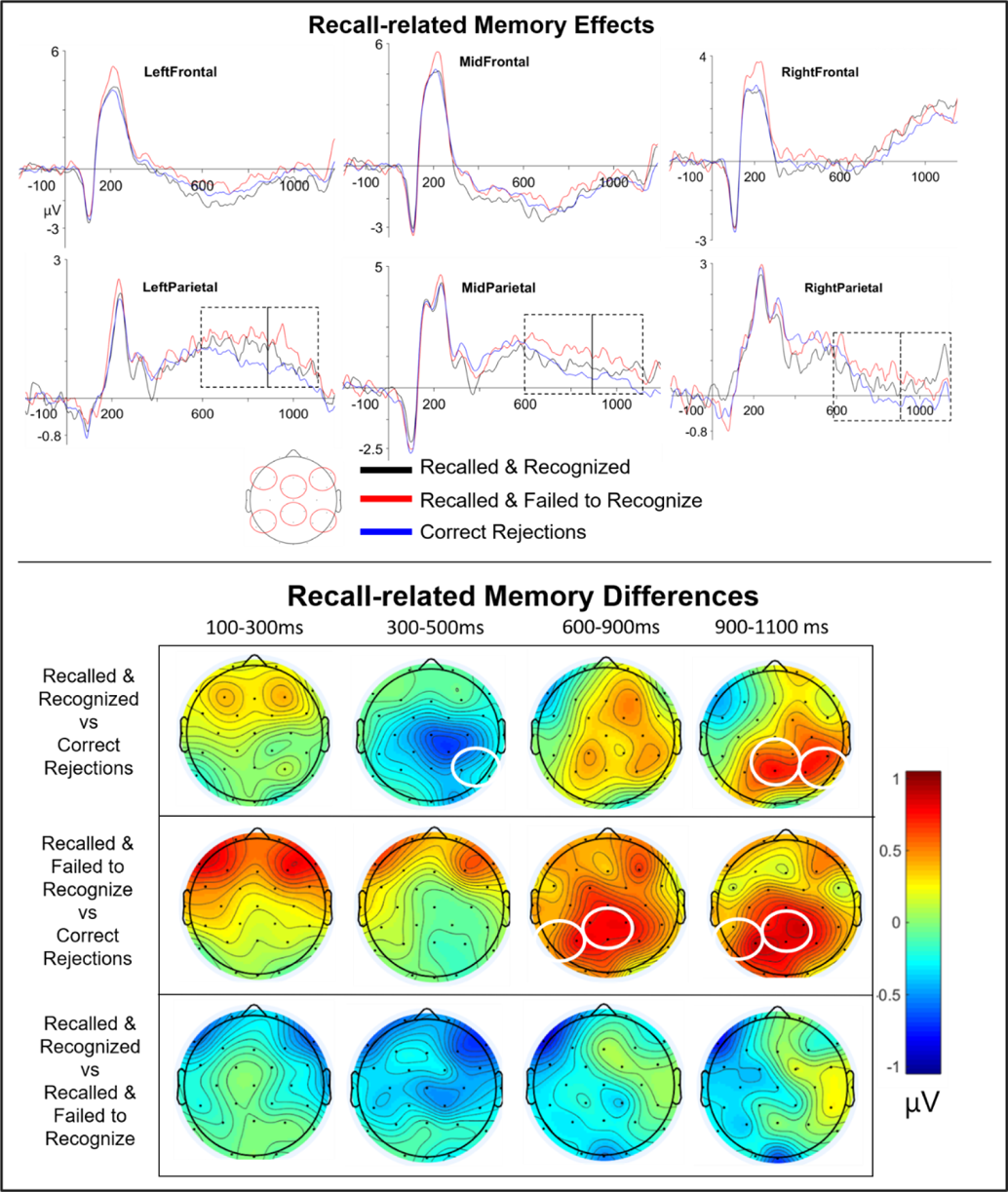
Recall-related Physiology. Top: ERPs of Recall Responses. Effects are shown for each of the six main electrode clusters analyzed, locations for which are illustrated in the representative topographic figure at the bottom. Dashed boxes indicate latencies which were found to exhibit significant effects at p <.05. Bottom: Topographic Maps of Recall Responses. Circles indicate where electrode clusters were found to be significantly different for each of the respective contrasts noted in the figure, below a threshold of p <.05.

Later, in the 600-900,ms latency, recognition failures were significantly more positive than correct rejections at left parietal sites (Misses: M = 1.30, SD = 2.44, CRs: M = .83, SD = 2.23, t(27) = 2.55, p < .05) and mid parietal sites (Misses: M = .46, SD = 3.26, CRs: M = −.17, SD = 3.04, t(27) = 2.55, p < .01). Then, in the 900-1100 ms latency, recognized recalls were significantly more positive than correct rejections at mid parietal electrodes (Hits: M = −.20, SD = 3.05, CRs: M = −.87, SD = 2.68, t(27) = 2.31, p < .05) and right parietal electrodes (Hits: M = −.37, SD = 2.53, CRs: M = −.82, SD = 2.30, t(27) = 2.16, p < .05). Recognition failures were also significantly more positive than correct rejections at left parietal sites (Misses: M = .93, SD = 2.16, CRs: M = .43, SD = 1.87, t(27) = 2.69, p < .01) and mid parietal sites (Misses: M = −.12, SD = 2.99, CRs: M = −.87, SD = 2.68, t(27) = 2.92, p < .01).

#### ERP Results: Recognition

We next sought to identify the ERP effects occurring during the ensuing latency when participants were answering the recognition prompt (i.e. whether the word that they had just produced was from the study phase, see Figure 3). The rationale for this analysis was that, since our conditions of interest (recall with recognition, and recall without recognition) were matched in recall success but varied in the recognition responses, it may be the neural activity occurring during the recognition responses (as opposed to that activity we first explore during recall responses) that determines which memory process is supporting the judgments; that is, we hypothesized that differences in behavior could have been due to activity occurring at recognition, as opposed to recall. We therefore examined the ERPs during the recognition epoch that immediately followed the time of the recall epoch for the same item (i.e., which occurred after participants had just seen the recall cue and made their recall response).

In the 100-300 ms recognition latency, at left frontal electrodes, recognized recalls (M = −1.70, SD = 2.96) were reliably more positive than recognition failures (M = −2.81, SD = 2.67), t(28) = 3.06, <u>p</u> < .01. Recognized recalls did not reliably differ from recognition failures or from correct rejections, as shown in Figure 6. Then, in the 300-500 ms recognition latency, at left frontal sites, recognized recalls (M = −2.20, SD = 3.99) were significantly more positive than correct rejections (M = −3.46, SD = 3.23), t(28) = 2.31, <u>p</u> < .05, and recognition failures (M = −4.17, SD = 3.46), t(28) = 3.12, <u>p</u> < .01. Recognized recalls (M = .19, SD = 2.50) were less positive than correct rejections (M = .90, SD = 1.89) at right parietal sites, t(28) = 2.52, <u>p</u> < .05. Recognition failures were also less positive than correct rejections both at left frontal sites (Misses: M = −4.17, SD = 3.46, CRs: M = −3.46, SD = 3.23, t(28) = 2.35, p < .05) and at right frontal sites (Misses: M = −4.01, SD = 4.04, CRs: M = −3.09, SD = 3.45, t(28) = 2.10, p < .05).

**Figure 6.**
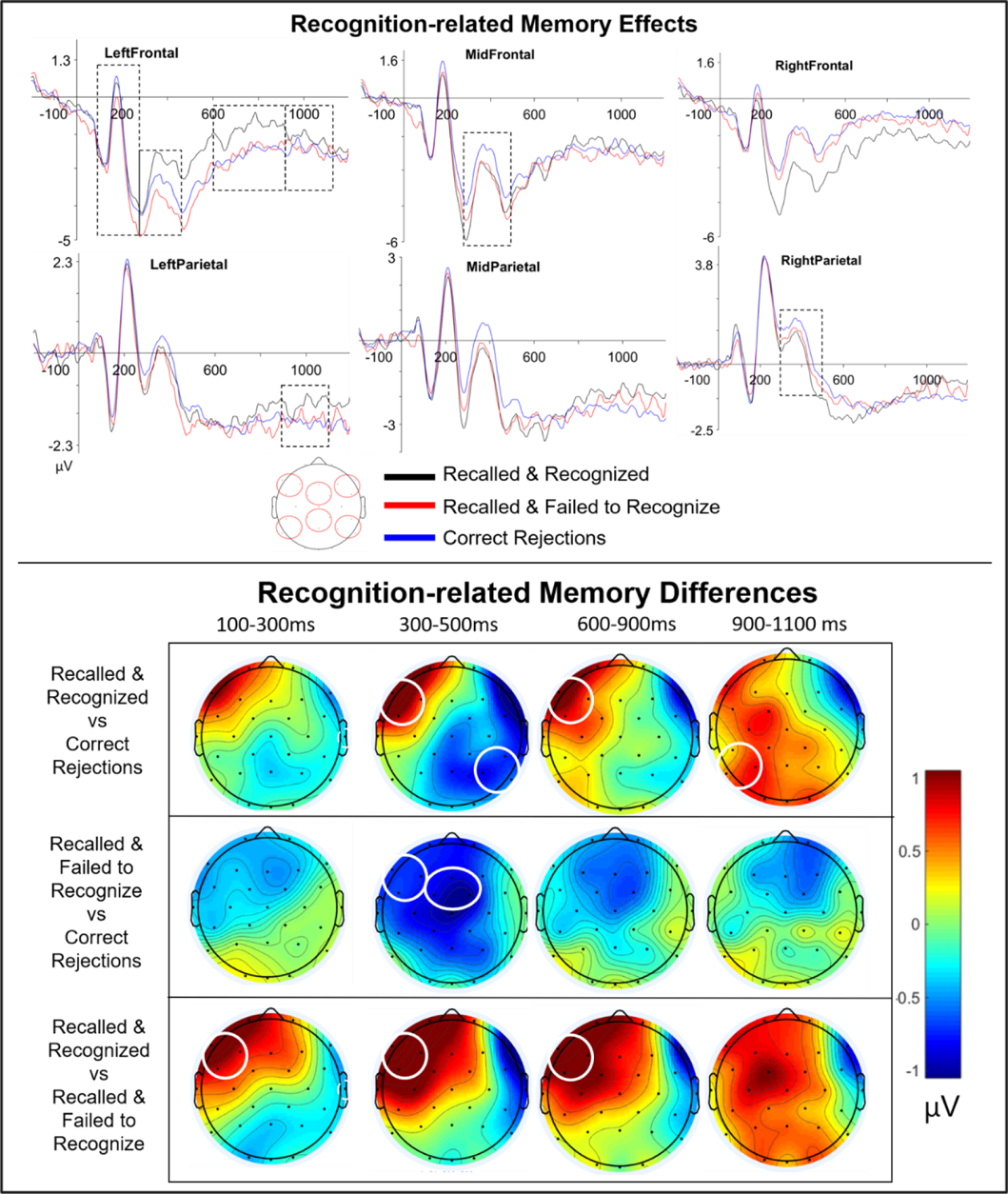
Figure 5: Recall-related Physiology. Top: ERPs of Recognition Responses. Effects are shown for each of the six main electrode clusters analyzed, locations for which are illustrated in the representative topographic figure at the bottom. Dashed boxes indicate latencies which were found to exhibit significant effects at p <.05. Bottom: Topographic Maps of Recognition Responses. Circles indicate where electrode clusters were found to be significantly different for each of the respective contrasts noted in the figure, below a threshold of p <.05.

During the 600-900 ms recognition latency, at left frontal sites, recognized recalls (M = −.70, SD = 3.64) were significantly more positive than correct rejections (M = −1.64, SD = 3.42), t(28) = 2.06, p < .05 and recognition failures (M = −2.01, SD = 3.42), t(28) = 2.46, p < .05. Then, during the 900-1100 ms recognition latency, at left parietal sites, recognized recalls (M = −1.16, SD = 2.53), were significantly more positive than correct rejections (M = −1.69, SD = 2.28), t (28 = 2.19, p < .05 (Figure 6). An overall summary of these ERP findings is presented as an integrated representation in Figure 7.

**Figure 7.**
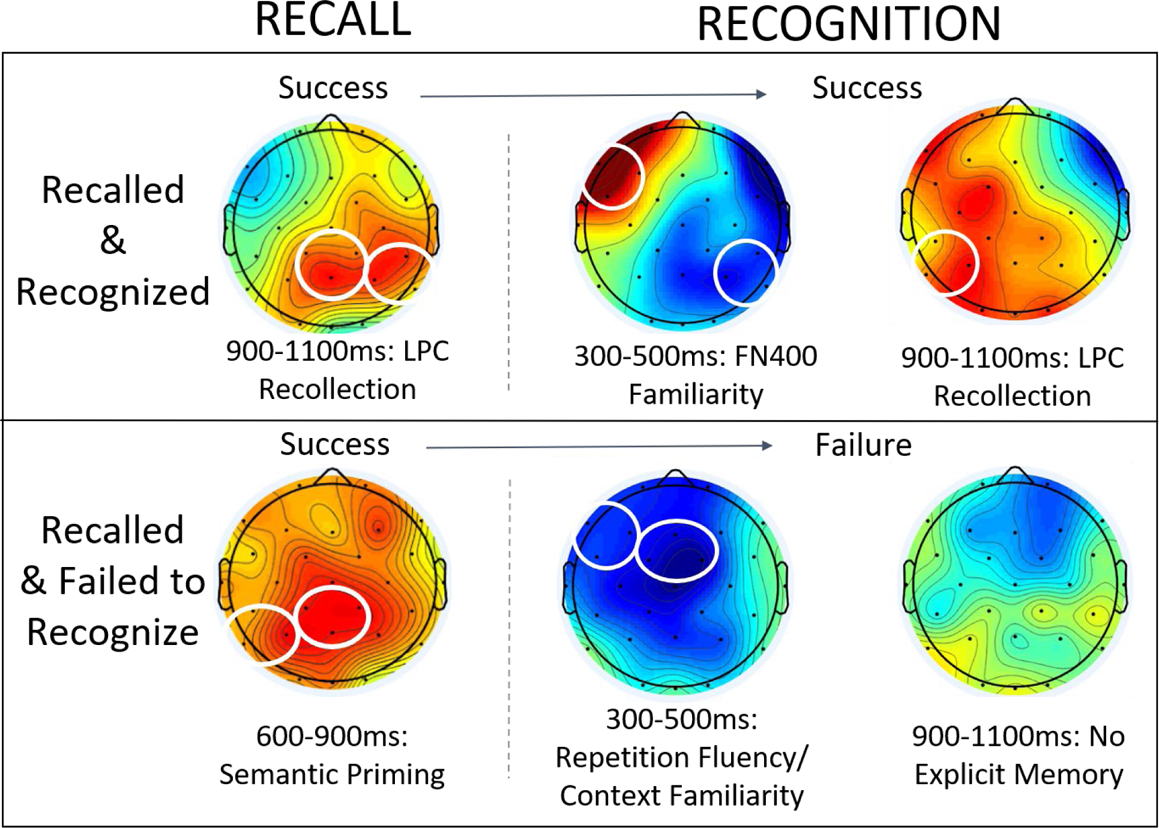
Summary Model of the ERP Data on Recall and Recognition Patterns. This illustrates the temporal sequence of activity as participants first process recall judgments for cued semantic associates, followed by old/new recognition judgments about the items that they just produced in the preceding recall response. Circles indicate where electrode clusters were found to be significantly different from correct rejections, below a threshold of p <.05.

## Discussion

In Experiment 2, we replicated the general pattern of behavioral findings exhibited in Experiments 1A and 1B (Figures 1 and 4), showing that recognition failures occur readily in the presence of cues. Our behavioral results further indicated faster reaction times for hits than for either misses or correct rejections at recognition, suggesting an ease of processing or “fluency” effect for true recalls. This finding converged with ERP results suggesting a role for fluency in supporting the recognition judgements of recognized recalls (e.g. Leynes & Addante, 2016), in addition to episodic memory effects of familiarity and recollection (Addante et al., 2012a,b; Leynes et al., 2005; for reviews, see Rugg & Curran, 2007). Recognized recalls represented processing emblematic of recollection, in that they exhibited effects which were slower responses followed by activity of familiarity early and recollection later in the epoch. In contrast to this, recognition failures exhibited processing reflecting a reliance on implicit priming in that the activity occurred fast and reflected ERP patterns consistent with what other studies have reported for fluency during early epochs which was distinct from familiarity-based processing, as discussed in further detail below.

One of the primarily goals of Experiment 2 was to better understand the processes that underlie the production of recognition failures in the first place: How could subjects produce words that they then could not recognize? In our main behavioral comparison of interest, recognition failure of recalled words was characterized by slower reaction times than successfully recognized recalled words when subjects made the recognition judgments of the combined response (see Figures 4). The differences between the reaction times in these conditions may represent a sequential search process in which participants search available memory for an old word and then, if they fail, they must think of a new word (Mecklinger, Rosburg, & Johannson, 2016). Although the behavioral studies of Experiment 1A and 1B did not measure response times for the recall response, Experiment 2 did so, which provided valuable insight into the processes recruited to produce such responses. At recall prompts, participants were faster to respond with an old word that was a correct cue-target pair than they were to respond with any old word that was from the study phase but not a pair, and also faster than they were to produce a new word.

Using ERPs, we found correlates of semantic priming for recognition failures of items that had been successfully recalled. When contrasting the physiological activity for the two epochs of interest (recall and recognition), overall, recall hits that went on to become recognition hits exhibited a different pattern of activity than did recall hits that went on to become recognitions misses, as described below. At recall, hits produced an LPC effect around 900-1100 ms (Figure 5), which suggests association with the putative neural correlates of recollection (Rugg & Curran, 2008). Then at recognition, this same condition of recalled hits was supported by early correlates of familiarity at left frontal sites that persisted through the epoch, followed by recollection-related LPC effects which emerged again at 900-1100 ms in parietal regions (Figure 5). Thus, recalled hits seem to be supported by a combination of both recollection and familiarity: first by recollection at recall epochs, and then sequentially by familiarity and recollection, respectively, that occurred thereafter during recognition epochs as the episodic information from the prior occurrence became further accessible to memory searches (e.g., Mecklinger et al., 2016).

On the other hand, the activity found during recall for recognition failures was notably different from that described for recognized recalls. At recall, a positive parietal effect appeared at 600-900 ms that might initially be thought to indicate recollection-related processing (Figure 5). However, upon further inspection of this effect’s features, a more logical explanation emerges to instead suggest it likely represents semantic priming. This is because the ERP effect is occurring earlier (600-900 ms) than the effects of recollection that were evident for the condition of recognized recalls (900-1100 ms), and the condition of being a recognition failure is not consistent with what would be expected with recollection. Furthermore, during recognition epochs, recognition failures were not associated with any of the explicit recognition-related physiology of recollection (Figure 6) that would have been expected to be consistent with a broad literature of recollection-related ERPs that occur during recognition memory (Addante, 2015; Addante, Ranganath, Olichney, & Yonelinas, 2012; Addante, Ranganath, & Yonelinas, 2012; Bridger et al., 2012; Li, Mao, Wang, & Guo, 2017; Rugg et al., 1998; Yu & Rugg, 2010; Bader & Mecklinger, 2017).

The preceding ERP analysis focused on EEG epochs for when participants were processing the cue word to make a recall response, but not upon EEG epochs from when participants were performing a recognition judgment about their recalled response. Upon reflection, we reasoned that the conditions being compared (recall hits that became recognition hits, and recall hits that became recognition misses) were each actually defined as the same conditions during recall, and that if we were seeking to understand why these behavioral conditions differed then we should also assess the physiology during the actual times in which the cognitive processes were happening that differentiated these conditions: specifically during recognition judgments. We hence conducted a further investigation of the recognition epochs, which revealed canonical physiological correlates of familiarity-based and recollection-based processing for conditions of recognized recalls^6^.

During the recognition latency, not only did recognition failures not exhibit ERP effects of explicit memory processes such as familiarity and recollection, but they were instead characterized by an early frontal negative-going effect that emerged around approximately 300-500 ms, and which was not present in the recognized recall condition (Figure 6). This effect is consistent with other left-frontal negative-going ERP reported of repetition fluency (Leynes & Zish, 2012; Leynes & Addante, 2016) and is similarly consistent with left-frontal negative-going ERP effects reported to occur a bit later in time (∼600 ms) for context familiarity (Addante, 2012a).

For ERPs during the recognition epoch of recognition failures, participants first experience semantic priming during initial recall—when they produce an old word from the study list—and then the word that they just produced is implicitly detected via repetition fluency or context familiarity (with the semantic nature being the familiar context) (Figure 7). Accordingly, the item is evidently lacking the conscious/explicit processing of item familiarity or recollection that would have been evident in an FN400 or LPC effect, respectively, like the one observed for recall with recognition items.

### General Discussion

Recognition failures in recall represent an unusual effect in episodic memory, in that participants are paradoxically able to recall words that they cannot then recognize as having been seen before. One would typically think that if an individual can successfully retrieve episodic information via recall that they would also then be able to recognize that information from the past; indeed, most memory models have traditionally assumed this, although there have been exceptions noted in the literature (Allan & Rugg, 1998; Angel, Fay, et al., 2010; Angel et al., 2009; Angel, Isingrini, et al., 2010; Rugg, Fletcher, et al., 1998). In Experiments 1A and 1B, we showed that just because an individual can “recall” a word does not mean that they actually remember it. Recognition failures occur readily in cued recall, regardless of whether they are detected with a recognition decision. Experiment 2 further elucidated why and how recognition failures occur. Most importantly, recognition failures appear to arise from fundamentally different processes than true recalls (i.e., recognized recalls). Whereas recognized recalls are initially produced from recollective processes, and then subsequently recognized via a combination of familiarity and recollection, recognition failures are initially produced from semantic priming, and show no evidence of traditional explicit memory processes at recognition (see Figure 7). Recognized recalls and recognition failures are thus of a qualitatively different kind. We turn now to a more detailed consideration of these conclusions.

#### The Impact of Cues and the Value of Recognition

The aim of the current investigation was to identify the neural and cognitive processes were that contribute to recognition failures. To achieve this, we adapted forced-recognition recall procedures to ERP, and also added a technological advance of integrating a digital voice-recorder that could time-stamp precisely when subjects experienced the phenomenon of memory recall, beyond behavioral methods traditionally used to measure recall. Together, these innovations each permitted us to successfully measure sensitive ERP conditions comparing both recall and recognition memory latencies sequentially (itself a relative innovation in ERP studies), which revealed distinct physiological effects that would not have been evident in the coarser analyses previously available in extant research.

A clear message from the convergent data across our experiments is that recognition failures are occurring in recall whether or not we measure them. When recognition is not measured after recall, cues appear to uniformly enhance memory; however, when recognition is measured, cues may not always be uniformly enhancing memory. We take note to also emphasize that the quality of the cues likely affects the relation, as weaker or stronger cues may lead to more or fewer recognition failures. Also, to be clear, it is not the case that cueing will never enhance memory per se; certainly, much research exists which can show the effectiveness of cues to aid memory retrieval. But, when cues are used, uncritically assuming that a correct recall from a retrieval cue represents explicit memory is a flawed assumption, as processes like implicit priming can clearly exist as alternate causes and contributory processes, and the ERP data from the current study reveal that the activation of implicit and explicit memory processes is occurring at different spatiotemporal profiles as discussed below. Moving forward, researchers should consider the factors that could contribute to a correct response on a recall trial in their tasks and, when possible, should include redundant measures to ensure that a “recall” really does represent explicit memory.

#### The Neural Correlates of Recall vs. Recognition

Recognized recalls displayed patterns of physiological activity different from those for recognition failures at recognition, suggesting that they derive from different cognitive processes. Recognition hits of recalled items were specifically characterized by early positive frontal activity, consistent with FN400 effects typically reported in the literature for familiarity-based processing (Friedman & Johnson, 2000; Rugg & Curran, 2007; Addante et al., 2012a). Such hits also showed late positive parietal activity that suggests an integration of recollection-based memory processing. Misses of recognition failures do not exhibit these same effects of recollection or familiarity, but instead exhibit a mid-frontal negative effect, which has been found by other studies to represent repetition fluency (Leynes & Zish, 2012; Leynes & Addante, 2016). It remains possible that repetition fluency was present for hits but was just overpowered by the stronger positive physiology for processes of familiarity and recollection that masked the negative-going effects of repetition fluency, but future research is needed to disentangle those features (e.g. Leynes & Addante, 2016).

One useful innovation of this project was the instrumentation of the analysis that separated semantic associate conditions. Because of this analysis, we were able to examine effects that would have otherwise been left undetected by traditional, coarser methods of analysis. Based on the data, we thus infer that combining conditions for non-associate and semantic associate words may dilute the priming effects that we observed, because in doing so, such a procedure is combining conditions that actually represent reliably different neurocognitive processing. As described earlier in Method, the traditional approach dating back to Tulving and Osler (1968) collapsed words which were produced from semantic associate cues and those from non-associate cues together into the same conditions: Counting items as successfully recalled regardless of which cue initiated their retrieval has been a frequently used practice in the recall literature (e.g.; Blaxton, 1989; Humphreys & Galbraith, 1975; Thomson & Tulving, 1970; Tulving & Osler, 1968). Not specifying conditions based on whether responses were generated from semantic associate cues or from non-associate cues may thus conflate cognitive processes if in fact distinct processes are used to arrive at these recalled items. In the current work, we reasoned that it may be possible to gain a more sensitive measure of our conditions of interest if we used an approach that instead separated the respective conditions to be more specific, based on whether a word was produced in response to a semantic associate versus to a non-associate cue. This approach revealed key differences and indicated that priming related to semantic associates was driving a core element of recalled items that went on to not be recognized from prior study. Separating recall and recognition responses based on whether they originated from semantic or non-associate cues may thus be an important consideration for future investigations of retrieval, too, and suggests that neuroimaging studies should inspect activity during both epochs of recall and recognition.

The current data suggests that the phenomenon of recognition failure of recalled items can be explained by a combination of implicit processes including semantic priming, repetition fluency (Leynes & Addante, 2016; Leynes & Zish, 2012), and potentially context familiarity (Addante et al., 2012a), such that failed recognition judgments of items successfully recalled do not evidently rely on familiarity or recollection, but instead evidently rely upon the activity associated with early semantic priming.

#### Adapting Recall for EEG

Overall, the current study makes several key contributions to the literature. First, it successfully demonstrates the ability to integrate voice-key technology into episodic memory paradigms to capture the precision of free recall responses, and to do so while concurrently recording electrophysiology from EEG. Second, it created several insights and innovations into the sensitivity of memory conditions for studying combined responses of recall and recognition in cued-recall paradigms. That is, we found that existing approaches in the field (i.e. Blaxton, 1989; Humphreys & Galbraith, 1975; Thomson & Tulving, 1970; Tulving & Osler, 1968) for measuring memory conditions can be successfully broken down into their constituent parts of distinct cognitive conditions, and that when this is done these parts were associated with distinct physiological patterns that would not otherwise have been detected. Third, the study also introduced the unusual step of analyzing and reporting sequential episodic memory epochs of both recall and recognition, identifying the differential patterns of neural activity occurring in each to support complex forms of memory retrieval.

Finally, these steps all converged to reveal novel insight into why and how recognized recalled items can be followed by recognition failures from recall of the same items. It appears that this phenomenon is due to cued semantic priming during recall followed by repetition fluency during recognition in the absence of explicit familiarity and recollection. Based upon the convergence of data here from our different experiments, we provide the insight that recall— long assumed to represent the exclusive domain of recollection-based processing in episodic memory (i.e. Yonelinas et al., 2002; Wais et al., 2004)—can at times also be served by a confluence of implicit cognitive processes including repetition fluency and semantic priming.

This suggests that future work can develop meaningful insights into these memory processes by using similar approaches that ensure the specificity of these response categories and measurement latencies. As such, the data are clear in suggesting two main conclusions: (1) that standard measures of cued recall can be contaminated by implicit memory, and (2) that treating cued recall responses as a relatively straightforward measure of explicit memory is not always appropriate.

The data & materials for experiments are available upon request; experiments were not pre-registered

## Acknowledgements

We thank the research assistants for their efforts in helping to collect the physiological data in Experiment 2: Alana Muller, Celene Gonzalez, Rose DeKock, Roman Lopez, Yoselin Canizales, Yesenia Casas, Kate Wright, and Summer Ledesma. We also thank Dionisio Amodeo and John Clapper for helpful comments on earlier drafts of that study’s results.

Experiment 1 was supported by a Natural Sciences and Engineering Research Council of Canada Discovery Grant (A7459) to CMM. Experiment 2 was supported by grants to RJA from the National Institute of Health (grant 1 L30 NS112849-01), the National Institute of Neurological Disorders and Stroke, the CSUSB Office of Academic Research Mini-Grants Program, the CSUSB Office of Student Research Faculty Assigned Time Grant Program, the CSUSB Office of Sponsored Programs Faculty Summer Research Fellowship, the CSUSB Faculty Senate Assigned Time for Exceptional Service to Students, and the CSUSB OSR Faculty-Student Research Grant to RJA & LAS.

LAS was supported in part by KBR’s Human Health and Performance Contract NNJ15HK11B through the National Aeronautics and Space Administration, as well as by the CSUSB Student Success Initiative’s Innovative Scholars Fund & Culminating Project Fellowships.

## Author Contributions

JDO designed and programmed Experiments 1A and 1B, collected and analyzed the data, and co-wrote the manuscript.

LAS programmed Experiment 2, collected Exp. 2 data, and analyzed Exp.2 data, and assisted with manuscript preparation.

FNA helped to collect the data for Exp. 1A and 1B.

CMM supervised the design and collection of data for Experiments 1A and 1B and contributed to the writing of the manuscript.

RJA designed Experiment 2, supervised the data collection, analyzed the data with LSA, and co-wrote the manuscript.

## Footnotes

1 For an interesting related phenomenon, see recognition-without-identification (RWI), a finding where participants have been shown to be able to recognize word-fragment cues of studied words even when they cannot complete the cue itself (i.e., they cannot ‘recall’ the word from a cue, if completing that cue is considered as recalling the item in the task; Ryal et al., 2011; Ryal & Cleary, 2012). In a way, RWI may represent the opposite of recognition failure (i.e.: where items are “recalled” in the absence of recognition).

2 As detailed by Paller, Lucas, and Voss (2012) and by Mecklinger et al. (2012), there has been some disagreement among researchers concerning the relative contributions of explicit and implicit processing to the mid-frontal ERP effect, particularly in studies using complex stimuli such as faces (Donaldson & Curran, 2007) or geometric shapes (Groh-Bordin, Zimmer, & Ecker, 2006; Voss et al., 2010). However, as described by Bridger et al., (2012), Mecklinger et al. (2012), and Yu and Rugg (2010), in most cases a reliable topographic dissociation can be observed, particularly when the stimuli are verbal.

3 This 200-ms delay in recall likely arises because, unlike in recognition tests where test items are presented at the onset of each test trial, in cued recall tests, participants are provided with a cue at the start of each test trial and must take a moment to generate their own candidate for recognition (see Generate-Recognize models of recall, e.g., Haist, Shimamura, & Squire, 1992; Jacoby & Hollingshead, 1990; Nobel & Shiffrin, 2001; Slamecka, 1972).

4 This version of correct rejections differs from the correct rejection condition examined by prior research because new words produced in response to semantic associate cues were not examined in this condition. We did not want to contaminate this “no memory” condition with a semantic associate cue, which could potentially have initiated any kind of memory process that was not subjectively reported by the subject.

5 Old items that were produced to cues other than their intended cue were rare (M = 5, SE = 0.51); these trials were excluded from the ERP analyses.

6 As previously mentioned, due to the more demanding nature of recall, we expected to find that physiological correlates of familiarity and recollection were delayed approximately 200 ms. However, for the recognition analysis, we did not expect the physiological effects of recollection and familiarity to be delayed because during this time period participants no longer had to produce a word, but instead simply had to rate the word that they already produced as being old or new.

## References

1. Addante, R. J. (2015). A critical role of the human hippocampus in an electrophysiological measure of implicit memory. Neuroimage, 109, 515–528.

2. Addante, R. J., Ranganath, C., Olichney, J., & Yonelinas, A. P. (2012). Neurophysiological evidence for a recollection impairment in amnesia patients that leaves familiarity intact. Neuropsychologia, 50(13), 3004–3014.

3. Addante, R. J., Ranganath, C., & Yonelinas, A. P. (2012). Examining ERP correlates of recognition memory: evidence of accurate source recognition without recollection. Neuroimage, 62(1), 439–450.

4. Addante, R. J., Watrous, A. J., Yonelinas, A. P., Ekstrom, A. D., & Ranganath, C. (2011). Prestimulus theta activity predicts correct source memory retrieval. Proceeding of the National Academy of Sciences USA, 108(26), 10702–10707.

5. Allan, K., Doyle, M. C., & Rugg, M. D. (1996). An event-related potential study of word-stem cued recall. Cognitive Brain Research, 4(4), 251–262.

6. Allan, K., & Rugg, M. D. (1998). Neural correlates of cued recall with and without retrieval of source memory. Neuroreport, 9(15), 3463–3466.

7. Allan, K., Wilding, E. L., & Rugg, M. D. (1998). Electrophysiological evidence for dissociable processes contributing to recollection. Acta Psychologica, 98(2-3), 231–252.

8. Angel, L., Fay, S., Bouazzaoui, B., Baudouin, A., & Isingrini, M. (2010). Protective role of educational level on episodic memory aging: an event-related potential study. Brain & Cognition, 74, 312–323.

9. Angel, L., Fay, S., Bouazzaoui, B., Granjon, L., & Isingrini, M. (2009). Neural correlates of cued recall in young and older adults: an event-related potential study. Neuroreport, 20(1), 75–79.

10. Angel, L., Isingrini, M., Bouazzaoui, B., Taconnat, L., Allan, K., Granjon, L., & Fay, S. (2010). The amount of retrieval support modulates age effects on episodic memory: Evidence from event-related potentials. Brain Research, 1335, 41–52.

11. Bader, R., & Mecklinger, A. (2017). Separating Event-related Potential Effects for Conceptual Fluency and Episodic Familiarity. Journal of Cognitive Neuroscience, 29(8), 1402–1414.

12. Bailey, C. H., Bartsch, D., & Kandel, E. R. (1996). Toward a Molecular Definition of Longterm Memory. Proceeding of the National Academy of Sciences USA, 93, 13445–13452.

13. Barco, A., Bailey, C. H., & Kandel, E. R. (2006). Common molecular mechanisms in explicit and implicit memory. Journal of Neurochemistry, 97(6), 1520–1533.

14. Bell, A. J., & Sejnowski, T. J. (1995). An information-maximization approach to blind separation and blind deconvolution. Neural Computation, 7(6), 1129–1159.

15. Berry, C. J., Shanks, D. R., & Henson, R. N. (2008). A single-system account of the relationship between priming, recognition, and fluency. Journal of Experimental Psychology Learning Mememory and Cognition, 34(1), 97–111.

16. Blaxton, T. A. (1992). Dissociations among memory measures in memory-impaired subjects: evidence for a processing account of memory. Memory and Cognition, 20(5), 549–562.

17. Bridger, E. K., Bader, R., Kriukova, O., Unger, K., & Mecklinger, A. (2012). The FN400 is functionally distinct from the N400. Neuroimage, 63(3), 1334–1342.

18. Bridger, E. K., Bader, R., & Mecklinger, A. (2014). More ways than one: ERPs reveal multiple familiarity signals in the word frequency mirror effect. Neuropsychologia, 57, 179–190.

19. Cohen, N. J., & Eichenbaum, H. (1993). Memory, amnesia, and the hippocampal system. Cambridge, MA: MIT Press.

20. Cohen, N. J., & Squire, L. R. (1980). Preserved learning and retention of pattern analyzing skill in amnesia: dissociation of knowing how and knowing that. Science, 210(4466), 207–2010.

21. Corkin, S. (2002). What’s new with the amnesic patient H.M.? Nature Reviews Neuroscience, 3, 153–160.

22. Curran, T. (2000). Brain potentials of recollection and familiarity. Memory & Cognition, 28(6), 923–938.

23. Curran, T., Tanaka, J. W., & Weiskopf, D. M. (2002). An electrophysiological comparison of visual categorization and recognition memory. *Cognitive, Affective*, & Behavioral Neurosci, 2(1), 1–18.

24. Delorme, A., & Makeig, S. (2004). EEGLAB: An open source toolbox for analysis of single-trial EEG dynamics including independent component analysis. Journal of Neuroscience Methods, 134(1), 9–21.

25. Diana, R. A., Yonelinas, A. P., & Ranganath, C. (2007). Imaging recollection and familiarity in the medial temporal lobe: A three-component model. Trends in Cognitive Science, 11(9), 379–386.

26. Donaldson, D. I., & Curran, T. (2007). Potential (ERP) studies of recognition memory for faces. Neuroimage, 36(2), 488–489.

27. Duzel, E., Yonelinas, A. P., Mangun, G. R., Heinze, H. J., & Tulving, E. (1997). Event-related brain potential correlates of two states of conscious awareness in memory. Proceedings of the National Academy of Sciences U S A, 94(11), 5973–5978.

28. Ezzyat, Y., Kragel, J. E., Burke, J. F., Levy, D. F., Lyalenko, A., Wanda, P., . . . Kahana, M. J. (2017). Direct brain stimulation modulates encoding states and memory performance in humans. Current Biology.

29. Fay, S., Isingrini, M., Ragot, R., & Pouthas, V. (2005). The effect of encoding manipulation on word-stem cued recall: An event-related potential study. Cognitive Brain Research, 24(3), 615–626.

30. Friedman, D., & Johnson, R., Jr. (2000). Event-related potential (ERP) studies of memory encoding and retrieval: A selective review. Microscopy Research and Technique, 51(1), 6–28.

31. Gabrieli, J. D. (1998). Cognitive neuroscience of human memory. Annual Reviews of Psychology, 49, 87–115.

32. Gardiner, J. M. (1988). Functional aspects of recollective experience. Memory and Cognition, 16, 309–311.

33. Gardiner, J. M., & Nilsson, L. G. (1993). Mathematical constraints and the Tulving-Wiseman law: A rejoinder. Memory, 1, 219–229.

34. Groh-Bordin, C., Zimmer, H. D., & Ecker, U. K. (2006). Has the butcher on the bus dyed his hair? When color changes modulate ERP correlates of familiarity and recollection. Neuroimage, 32(4), 1879–1890.

35. Gruber, M. J., & Otten, L. J. (2010). Voluntary control over prestimulus activity related to encoding. Journal of Neuroscience, 30(29), 9793–9800.

36. Haist, F., Shimamura, A. P., & Squire, L. R. (1992). On the relationship between recall and recognition. Journal of Experimental Psychology: Learning Memory and Cognition, 18(4), 691–702.

37. Hannula, D. E., & Greene, A. J. (2012). The hippocampus reevaluated in unconscious learning and memory: At a tipping point? Frontiers in Human Neuroscience, 6, 80.

38. Jacoby, L. L. (1991). Separating automatic from intentional uses of memory. Journal of Memory and Language, 30, 513–541.

39. Jacoby, L. L., & Hollingshead, A. (1990). Toward a generate/recognize model of performance on direct and indirect tests of memory. Journal of Memory and Language, 29(4), 433–454.

40. Jernigan, T. L., Ostergaard, A. L., & Fennema-Notestine, C. (2001). Mesial temporal, diencephalic, and striatal contributions to deficits in single word reading, word priming, and recognition memory. Journal of the International Neuropsycholigcal Society, 7(1), 63–78.

41. Kahana, M. J. (2006). The cognitive correlates of human brain oscillations. Journal of Neuroscience, 26(6), 1669–1672.

42. Kim, A.S., Vallesi, A., Picton, T.W., Tulving, E., 2009. Cognitive association formation in episodic memory: Evidence from event-related potentials. Neuropsychologia, 47(14), 3162–3173.

43. Kintsch, W. (1968). Recognition and free recall of organized lists. Journal of Experimental Psychology, 78(3), 481–487.

44. Kintsch, W. (1970). Models for free recall and recognition. In D. A. Norman (Ed.), Models of human memory (pp. 331–373). New York: Academic Press.

45. Klem, G. H., Luders, H. O., Jasper, H. H., & Elger, C. (1999). The ten–twenty electrode system of the international federation. The international federation of clinical neurophysiology. Electroencephalography and Clinical Neurophysiological Supply, 52, 3–6.

46. Kragel, J. E., Ezzyat, Y., Sperling, M. R., Gorniak, R., Worrell, G. A., Berry, B. M., . . . Kahana, M. J. (2017). Similar patterns of neural activity predict memory function during encoding and retrieval. Neuroimage, 155, 60–71.

47. Leynes, A., & Zish, K. (2012). Event-related potential (ERP) evidence for fluency-based recognition memory. Neuropsychologia, 50(14), 3240–3249.

48. Leynes, P. A., & Addante, R. J. (2016). Neurophysiological evidence that perceptions of fluency produce mere exposure effects. *Cognitive, Affective*, & Behavioral Neuroscience, 16(4), 754–767.

49. Leynes, P. A., Landau, J., Walker, J., & Addante, R. J. (2005). Event-related potential evidence for multiple causes of the revelation effect. Consciousness & Cognition, 14(2), 327–350.

50. Li, B., Mao, X., Wang, Y., & Guo, C. (2017). Electrophysiological correlates of familiarity and recollection in associative recognition: Contributions of perceptual and conceptual processing to unitization. Frontiers in Human Neuroscience, 11, 125.

51. Li, B., Taylor, J. R., Wang, W., Gao, C., & Guo, C. (2017). Electrophysiological signals associated with fluency of different levels of processing reveal multiple contributions to recognition memory. Consciousness & Cognition, 53, 1–13.

52. Lockhart, R. S., Craik, F. I., & Jacoby, L. (1976). Depth of processing, recognition and recall. In J. Brown (Ed.), Recall and recognition. Oxford, UK: John Wiley & Sons.

53. Lopez-Calderon, J., & Luck, S. J. (2014). ERPLAB: An open-source toolbox for the analysis of event-related potentials. Frontiers in Human Neuroscience, 8, 213.

54. Lucas, H. D., Paller, K. A., & Voss, J. L. (2012). On the pervasive influences of implicit memory. Cognitive Neuroscience, 3(3-4), 219–226.

55. Luck, S. J. (2014). An introduction to the Event-Related Potential technique (2nd ed.). Cambridge, MA: MIT Press.

56. Mandler, G. (1980). Recognizing: The judgment of previous occurrence. Psychological Review, 87(252-271).

57. Martin, E. (1975). Generation-recognition theory and the encoding specificity principle. Psychological Review, 82(2), 150–153.

58. Mecklinger, A. (2006). Electrophysiological measures of familiarity memory. Clinical EEG Neurosciencei, 37(4), 292–299.

59. Mecklinger, A., Frings, C., & Rosburg, T. (2012). Response to Paller et al.: The role of familiarity in making inferences about unknown quantities. Trends in Cognitive Science, 16(6), 315–316.

60. Nelson, D. L., McEvoy, C. L., & Schreiber, T. A. (2004). The University of South Florida free association, rhyme, and word fragment norms. Behavior Research Methods, Instruments, & Computers, 36(3), 402–407.

61. Nobel, P. A., & Shiffrin, R. M. (2001). Retrieval processes in recognition and cued recall. Journal of Experimental Psychology: Learning, Memory, and Cognition, 27(2), 384–413.

62. Nomi, J. S., & Cleary, A. M. (2012). Judgments for inaccessible targets: Comparing recognition without identification and the feeling of knowing. Memory & Cognition, 40(8), 1178–1188.

63. Oldfield, RC (March 1971). The assessment and analysis of handedness: The Edinburgh inventory. Neuropsychologia. 9 (1): 97–113.

64. Ostergaard, A. L. (1999). Priming deficits in amnesia: Now you see them, now you don’t. Journal of the International Neuropsychological Society, 5(3), 175–190.

65. Otten, L.J., Quayle, A.H., Akram, S., Ditewig, T.A., Rugg, M.D., 2006. Brain activity before an event predicts later recollection. Nature Neuroscience., 9 \(4), 489–491.

66. Paller, K. A., Lucas, H. D., & Voss, J. L. (2012). Assuming too much from ‘familiar’ brain potentials. Trends in Cognitive Science, 16(6), 313–315; discussion 315-316.

67. Polster, M. R., Nadel, L., & Schacter, D. L. (1991). Cognitive neuroscience analyses of memory: A historical perspective. Journal of Cognitive Neuroscience, 3(2), 95–116.

68. Ramos, T., Marques, J., & Garcia-Marques, L. (2017). The memory of what we do not recall: Dissociations and theoretical debates in the study of implicit memory. Psicológica, 38, 365–393.

69. Reagh, Z. M., & Ranganath, C. (2018). What does the functional organization of cortio-hippocampal networks tell us about the functional organization of memory? Neuroscience Letters, volume 680, 69–76, https://doi.org/10.1016/j.neulet.2018.04.050.

70. Reber, P. J. (2013). The neural basis of implicit learning and memory: A review of neuropsychological and neuroimaging research. Neuropsychologia, 51(10), 2026–2042.

71. Rosenthal, C. R., & Soto, D. (2016). The anatomy of non-conscious recognition memory. Trends in Neuroscience, 39(11), 707–711.

72. Rugg, M. D., & Curran, T. (2007). Event-related potentials and recognition memory. Trends in Cognitive Science, 11(6), 251–257.

73. Rugg, M. D., Fletcher, P. C., Allan, K., Frith, C. D., Frackowiak, R. S., & Dolan, R. J. (1998). Neural correlates of memory retrieval during recognition memory and cued recall. Neuroimage, 8(3), 262–273.

74. Rugg, M. D., Mark, R. E., Walla, P., Schloerscheidt, A. M., Birch, C. S., & Allan, K. (1998). Dissociation of the neural correlates of implicit and explicit memory. Nature, 392(6676), 595–598.

75. Rugg, M. D., Walla, P., Schloerscheidt, A. M., Fletcher, P. C., Frith, C. D., & Dolan, R. J. (1998). Neural correlates of depth of processing effects on recollection: Evidence from brain potentials and positron emission tomography. Experimental Brain Research, 123(1-2), 18–23.

76. Ryals, A. J., Yadon, C. A., Nomi, J. S., & Cleary, A. M. (2011). When word identification fails: ERP correlates of recognition without identification and of word identification failure. Neuropsychologia, 49(12), 3224–3237.

77. Schacter, D., Chui, C. Y. P., & Ochsner, K. N. (1993). Implicit memory: A selective review. Annu Reviews of Neuroscience, 16, 159–182.

78. Slamecka, N. J. (1972). The question of associative growth in the learning of categorized material. Journal of verbal learning and verbal behavior, 11(3), 324–332.

79. Squire, L. R. (2004). Memory systems of the brain: A brief history and current perspective. Neurobiology of Learning and Mememory, 82(3), 171–177.

80. Squire, L. R., & Dede, A. J. (2015). Conscious and unconscious memory systems. Cold Spring Harbor Perspectives in Biology, 7(3), a021667.

81. Squire, L. R., & Zola-Morgan, S. (1991). The medial temporal lobe memory system. Science, 253, 1380–1386.

82. Strozak, P., Abedzadeh, D., & Curran, T. (2016). Separating the FN400 and N400 potentials across recognition memory experiments. Brain Research, 1635, 41–60.

83. Suthana, N., & Fried, I. (2012). Percepts to recollections: Insights from single neuron recordings in the human brain. Trends Cogn Sci, 16(8), 427–436.

84. Thomson, D. M., & Tulving, E. (1970). Associative encoding and retrieval: Weak and strong cues. Journal of Experimental Psychology, 86(2), 255–262.

85. Tulving, E. (1985). How many memory systems are there? American Psychologist. Vol 40(4), 385–398.

86. Tulving, E. (1993). What is episodic memory? Current Directions in Psychological Science. Vol. 2 (3), 67–70

87. Tulving, E., & Markowitsch, H. J. (1998). Episodic and declarative memory: Role of the hippocampus. Hippocampus, 8(3), 198–204.

88. Tulving, E., & Osler, S. (1968). Effectiveness of retrieval cues in memory for words. Journal of Experimental Psychology, 77(4), 593–601.

89. Tulving, E., & Schacter, D. (1990). Priming and human memory systems. Science, 247, 301–305.

90. Tulving, E., & Thomson, D. M. (1973). Encoding specificity and retrieval processes in episodic memory. Psychological Review, 80(5), 352–373.

91. Tulving, E., & Watkins, O. C. (1977). Recognition failure of words with a single meaning. Memory & Cognition, 5(5), 513–522.

92. Voss, J. L., Lucas, H. D., & Paller, K. A. (2010). Conceptual priming and familiarity: Different expressions of memory during recognition testing with distinct neurophysiological correlates. Journal of Cognitive Neuroscience, 22(11), 2638–2651.

93. Voss, J. L., Lucas, H. D., & Paller, K. A. (2012). More than a feeling: Pervasive influences of memory without awareness of retrieval. Cognitive Neuroscience, 3(3-4), 193–207.

94. Voss, J. L., & Paller, K. A. (2007). Neural correlates of conceptual implicit memory and their contamination of putative neural correlates of explicit memory. Learning & Memory, 14(4), 259–267.

95. Voss, J. L., & Paller, K. A. (2017). Neural Substrates of Remembering – Electroencephalographic Studies, In Learning and Memory: A Comprehensive Reference, edited by John H. Byrne, Academic Press, Oxford, 2008, Pages 79–97.

96. Wilding, E. L. (2006). The practice of rescaling scalp-recorded event-related potentials. Biological Psychology, 72(3), 325–332.

97. Wolk, D. A., Schacter, D. L., Berman, A. R., Holcomb, P. J., Daffner, K. R., & Budson, A. E. (2004). An electrophysiological investigation of the relationship between conceptual fluency and familiarity. Neuroscience Letters, 369(2), 150–155.

98. Woodruff, C. C., Hayama, H. R., & Rugg, M. D. (2006). Electrophysiological dissociation of the neural correlates of recollection and familiarity. Brain Research, 1100(1), 125–135.

99. Yassa, M. A., & Stark, C. E. (2011). Pattern separation in the hippocampus. Trends in Neuroscience, 34(10), 515–525.

100. Yonelinas, A. P. (1994). Receiver-operating characteristics in recognition memory: Evidence for a dual-process model. *Journal of Experimental Psychology: Learning*, Memory, and Cognition, 20(6), 1341–1354.

101. Yonelinas, A. P. (2001). Components of episodic memory: The contribution of recollection and familiarity. Philosophical Transactions of the Royal Society of London Biological Sciences, 356(1413), 1363–1374.

102. Yonelinas, A. P. (2002). The nature of recollection and familiarity: A review of 30 years of research. Journal of Memory and Language, 46(3), 441–517.

103. Yonelinas, A. P., Aly, M., Wang, W. C., & Koen, J. D. (2010). Recollection and familiarity: Examining controversial assumptions and new directions. Hippocampus, 20(11), 1178–1194.

104. Yonelinas, A. P., Kroll, N. E., Quamme, J. R., Lazzara, M. M., Sauve, M. J., Widaman, K. F., & Knight, R. T. (2002). Effects of extensive temporal lobe damage or mild hypoxia on recollection and familiarity. Nature Neuroscience, 5(11), 1236–1241.

105. Yonelinas, A. P., Quamme, J. R., Widaman, K. F., Kroll, N. E., Sauve, M. J., & Knight, R. T. (2004). Mild hypoxia disrupts recollection, not familiarity. *Cognitive*, Affective, and Behavioral Neuroscience, 4(3), 393–400; discussion 401-406.

106. Yu, S. S., & Rugg, M. D. (2010). Dissociation of the electrophysiological correlates of familiarity strength and item repetition. Brain Research, 1320, 74–84.

